# nf-cavalier: A Nextflow Pipeline for Rare Disease Variant Prioritization and Reporting

**DOI:** 10.64898/2026.08.06.743410

**Authors:** Jacob E. Munro, Joshua Reid, Melanie Bahlo, Mark F. Bennett

## Abstract

**nf-cavalier** is a Nextflow pipeline that automates genomic variant annotation, filtering, and reporting for individuals with rare Mendelian diseases. The pipeline takes as input variant callsets for an individual, family, or rare disease cohort, together with a target gene panel or a phenotype of interest. Variants are then filtered using various customisable criteria, including predicted gene consequence, computational pathogenicity predictions, population frequency, and familial segregation. The sequencing data for candidate variants is then visualised for human review. Candidate variant results are returned in user-friendly output formats, including interactive HTML reports and PowerPoint slide decks, with embedded links to external resources that enable rapid review by clinical research teams. **nf-cavalier** is maintained on GitHub (bahlolab/nf-cavalier) and licensed under the permissive MIT open-source licence.

## Statement of need

Identifying candidate pathogenic genetic variants from genome sequencing data is a complex task that requires coordinating a wide range of software and data sources. Millions of variant calls are generated per individual and must be interpreted alongside known disease gene associations (e.g., PanelApp^1^, HPO^2^), gene and transcript models (e.g., GENCODE^3^), computational pathogenicity predictions (e.g., CADD^4^), population frequency data (e.g., gnomAD^5^) and known pathogenic variant databases (e.g., ClinVar^6^). Variant inheritance patterns must be considered in familial cases, with population frequency filters tailored to dominant, recessive, and compound heterozygous inheritance patterns. Finally, candidate variants must be reported, and the sequencing data supporting the variant visualised for human review prior to further validation, segregation, and pathogenicity classification.

This task is further complicated by the need to jointly interpret short variants (single-nucleotide variants (SNVs; 1 bp) and insertion-deletion variants (indels; 2 – 50 bp)) and structural variants (larger alterations; > 50 bp), which are typically identified using separate workflows.

Several software tools are commonly used for the individual tasks carried out in this workflow. For example, VEP^7^ handles variant annotations and consequence prediction based on specified gene models, but does not support inheritance-based filtering or variant visualisation. slivar^8^ enables rapid querying of variant call format (VCF) files with an expressive syntax that allows specific inheritance models to be interrogated; however, it does not perform annotation or visualisation. Tools such as igv-reports^9^ and SVPV^10^ generate visualisations of sequencing data to aid variant review, but do not perform prioritisation. As such, there is a need for tools that orchestrate and coordinate a variety of task-specific applications into a unified variant prioritisation and reporting framework, improving both reproducibility and user-friendliness. Such integrated workflows are easier to maintain, apply consistently, and validate than fragmented, ad hoc approaches.

**nf-cavalier** automates this process in a flexible, extensible, and reproducible manner, generating user-friendly outputs in interactive HTML and PowerPoint formats that can be shared and reviewed by multiple collaborators. Metadata describing gene lists and runtime parameters are embedded in the HTML reports, enabling previous analyses to be audited and reproduced. By integrating annotation, inheritance-based filtering, visualisation, and reporting into a single workflow, **nf-cavalier** reduces the manual effort required for rare disease variant prioritisation while improving reproducibility. The target audience for **nf-cavalier** is clinical research teams exploring the genetic causes of rare diseases.

## State of the field

There are several comparable tools for rare disease variant prioritisation; most notably seqr^12^, Scout^13^, and MOLGENIS-VIP^11^ (Table 1). Of these, VIP is the closest comparator to **nf-cavalier**, as both tools are Nextflow-based and produce static HTML reports without requiring a server or database backend. This makes them suitable for institutional HPC environments where persistent server infrastructure is typically unsupported or undesirable. In contrast, seqr and Scout depend on continuously running web services, which makes them out of reach for many research teams due to ongoing costs and infrastructure requirements. The recommended seqr deployment uses Kubernetes to orchestrate multiple interdependent services, including SQL databases (PostgreSQL, ClickHouse), Redis (cache), the seqr application, and a pipeline-runner service for loading variants into the databases. Scout is simpler, with the recommended deployment using two Docker services: MongoDB and the Scout web application. Many institutional HPC environments restrict or do not support Kubernetes and Docker deployments.

**Table 1.**
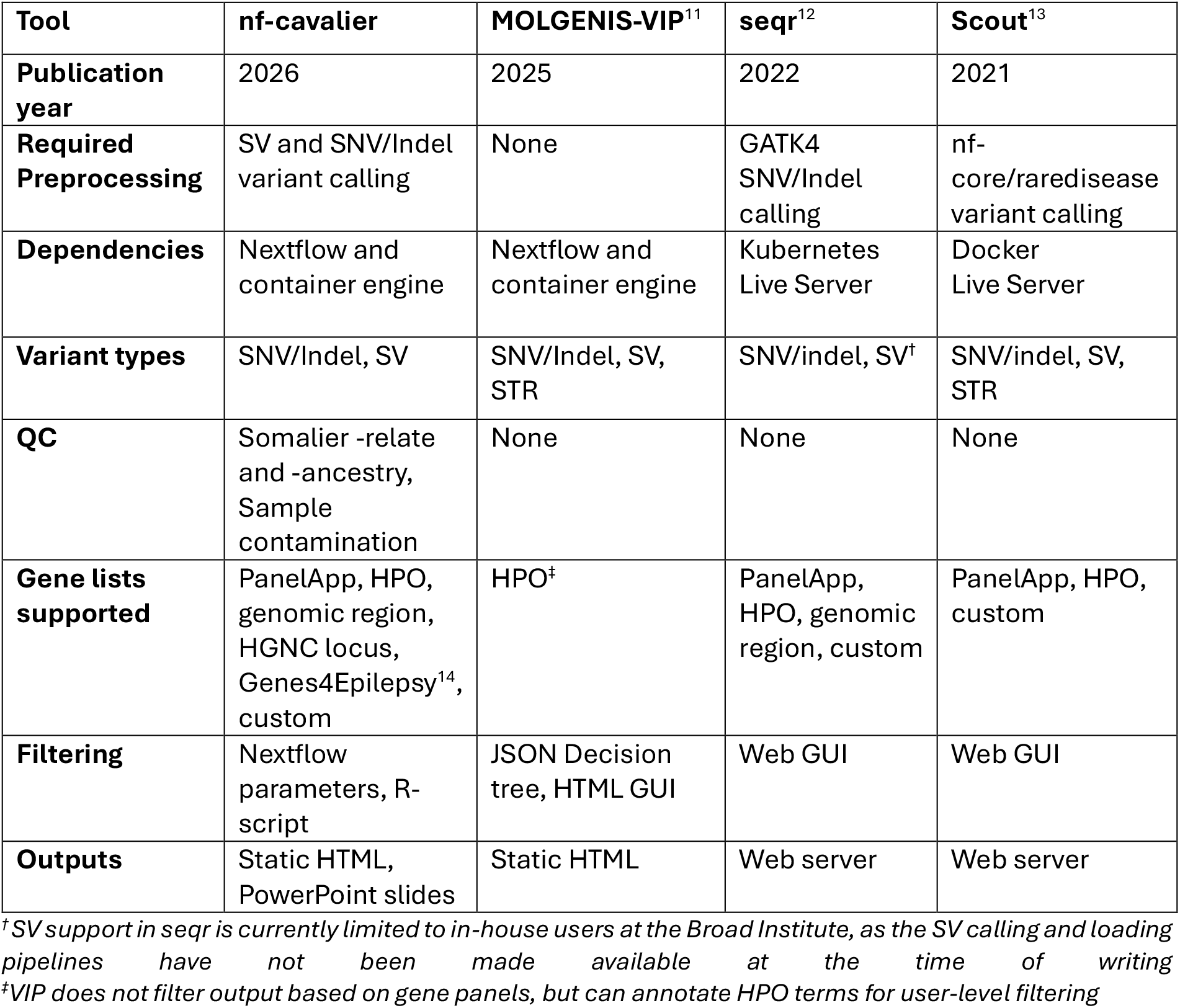
Comparison of **nf-cavalier** to leading variant prioritisation tools.

**nf-cavalier** and VIP are designed for different use cases. VIP targets production-oriented clinical or diagnostic variant classification, while **nf-cavalier** targets exploratory, tunable research candidate discovery and prioritisation. VIP encodes interpretation rules as a configurable decision tree in JSON, combining evidence criteria to assign variants to pathogenicity classifications based on ACMG guidelines. In contrast, **nf-cavalier** does not attempt automated classification; instead, it applies configurable filtering criteria to identify candidate variants for downstream review and interpretation. These criteria, including allele frequency, genotype quality, consequence class, and gene-list membership, are exposed as independently configurable, named parameters. For example, `FILTER_SHORT_POP_DOM_MAX_AF = 0.0001` sets the maximum gnomAD allele frequency allowed for dominantly inherited short variants. This provides a transparent representation of the filtering strategy, allowing researchers to inspect, understand, and iteratively refine individual filtering criteria in a straightforward manner.

## Software design

Variant prioritisation pipelines are subject to substantial churn – new annotation sources become available, new pathogenicity-prediction methods are released, and new classes of variation are recognised. **nf-cavalier** is designed to allow new annotation datasets, pathogenicity predictions, and population frequency resources to be incorporated with minimal effort, and the framework is left open to incorporating new approaches as they are released. The pipeline is fundamentally anchored in the VCF specification, allowing compatibility with a range of upstream variant callers and annotation resources. This makes **nf-cavalier** particularly suitable for research pipelines, which are often at the cutting edge of new variant discovery.

The pipeline has been implemented in Nextflow, leveraging Nextflow’s framework for reproducibility, containerisation, and ability to run on local machines, high-performance computing clusters, and cloud environments. The pipeline consists of three modules: annotation, QC, and reporting (Figure 1). We also include a one-time setup pipeline that automates the download of required reference datasets, as well as a script to download a demonstration dataset based on the 1000 Genomes CEPH trio for testing and evaluation.

**Figure 1.**
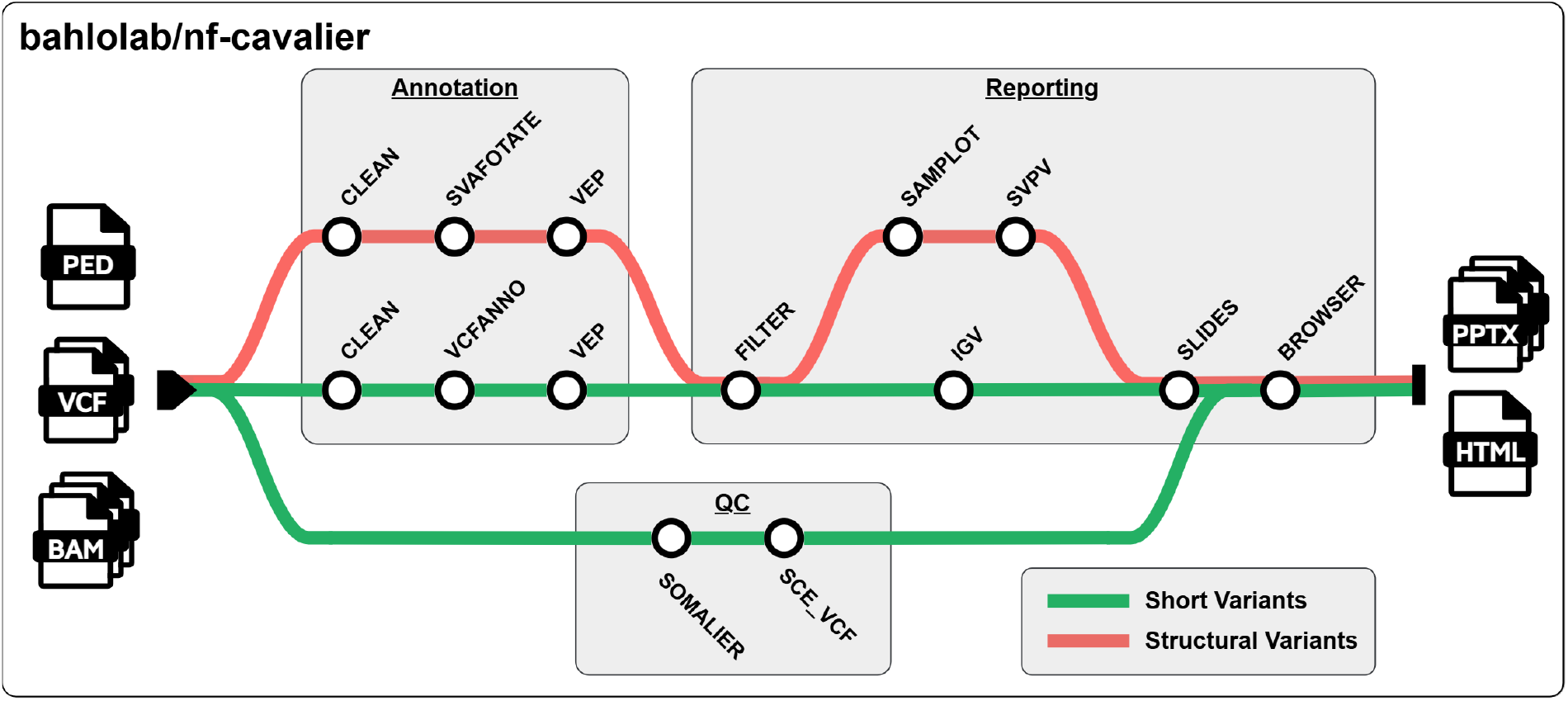
Schematic overview of **nf-cavalier** pipeline

Annotation takes cohort VCFs (short and/or structural variants) as input, splits them into shards for parallel processing, and performs variant normalisation and annotation with standard tools bcftools^15^, vcfanno^16^, VEP^7^, and SVAFotate^17^. Sharding improves scalability, reducing runtime from days to hours depending on available compute resources. vcfanno is employed to rapidly incorporate pre-calculated variant-specific metrics for short variants, including population allele frequencies from gnomAD v4.1, known pathogenicity classifications from ClinVar, predicted deleteriousness scores from CADD, as well as arbitrary user-specified annotation sources. VEP is used to add gene-level consequence predictions as well as annotations that depend on transcript models, including REVEL^18^, AlphaMissense^19^, and SpliceAI^20^ scores. For structural variants, SVAFotate is used to transfer gnomAD v4.1 SV frequencies using an overlap-based matching approach.

Filtering operates on a per-family basis and applies user-configurable filters for genes of interest (derived from PanelApp, HPO, genomic regions, or custom inputs), genic consequence (e.g. VEP impact or consequences), and deleteriousness predictions. By default, all reported ClinVar pathogenic/likely pathogenic variants in a gene of interest are retained, and all reported ClinVar benign or likely benign variants are excluded. Variants are then matched to inheritance models – all affected family members must carry dominant variants; recessive variants must be present only in a homozygous state in affected family members; and compound heterozygous pairs must be present together only in affected family members. Allele frequency filters are then applied depending on inheritance model, with the default dominant threshold set at 1/10,000 and the default recessive threshold set at 1/100.

The reporting stage visualises candidate variants using igv-reports, SVPV, or Samplot to support manual review and quality control. Reports are compiled into PowerPoint or PDF slide decks (Figure 2) and published alongside a static HTML report that enables interactive querying and exploration of variants (Figure 3). The HTML report also provides live access to other resources via public APIs, including PanelApp, HPO, ClinVar, gnomAD, and the UCSC genome browser.

**Figure 2.**
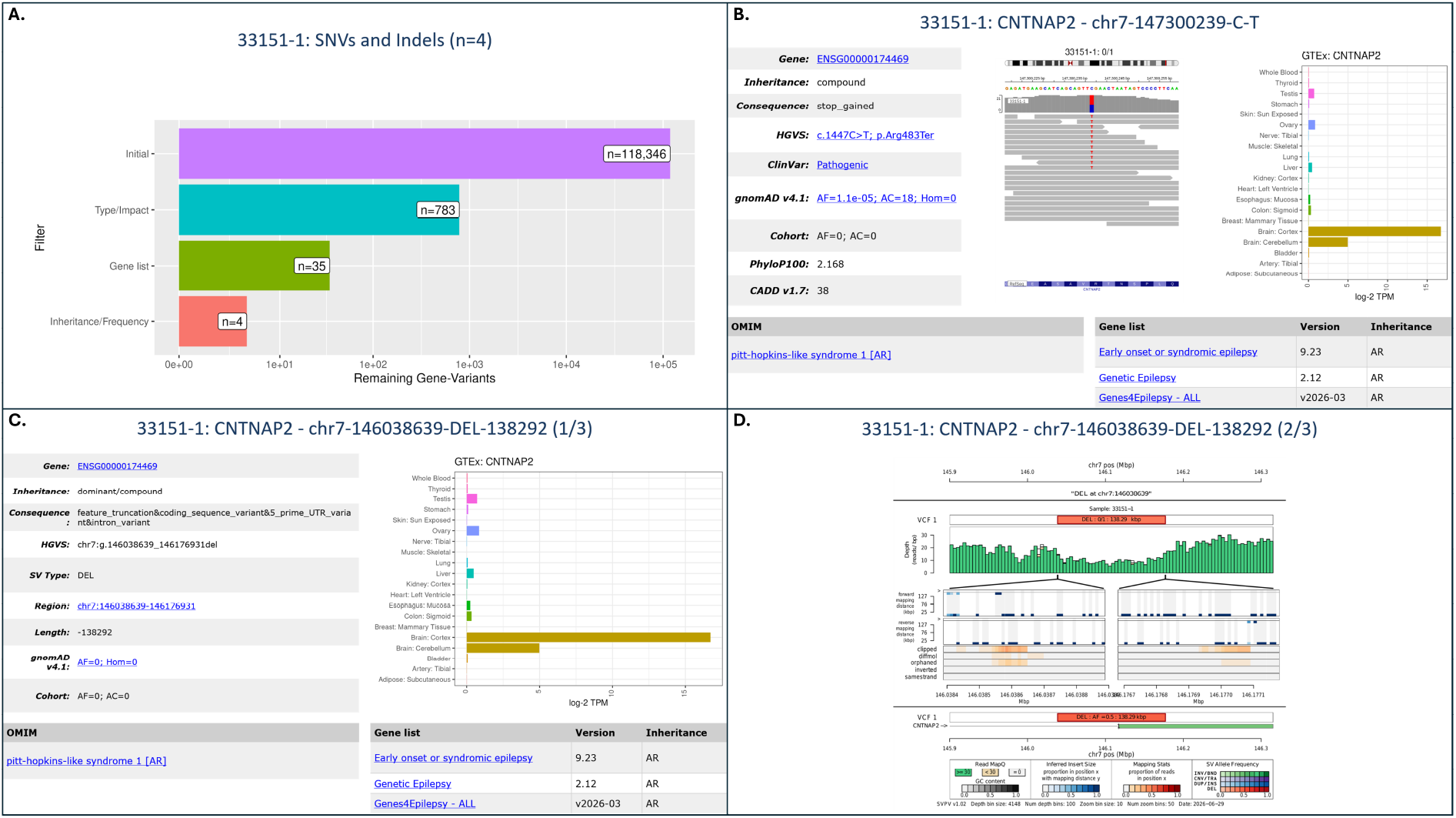
Example **nf-cavalier** variant PowerPoint output slides for a research participant with compound heterozygous pathogenic variants in the gene CNTNAP2. A) Summary slide detailing counts after each level of SNV/Indel filtering was applied, resulting in four candidate variants for this example analysis. B. PowerPoint output slide with details on a stop-gain variant in CNTNAP2, known to be pathogenic in ClinVar. Blue text represents hyperlinks to external resources. C. PowerPoint output slide detailing a 138 kb deletion affecting CNTNAP2, flagged by **nf-cavalier** as potentially compound heterozygous. D. nf-cavalier PowerPoint slide embedding an SVPV plot of the structural variant for inspection and QC. This case has been published in Munro et al^21^.

**Figure 3.**
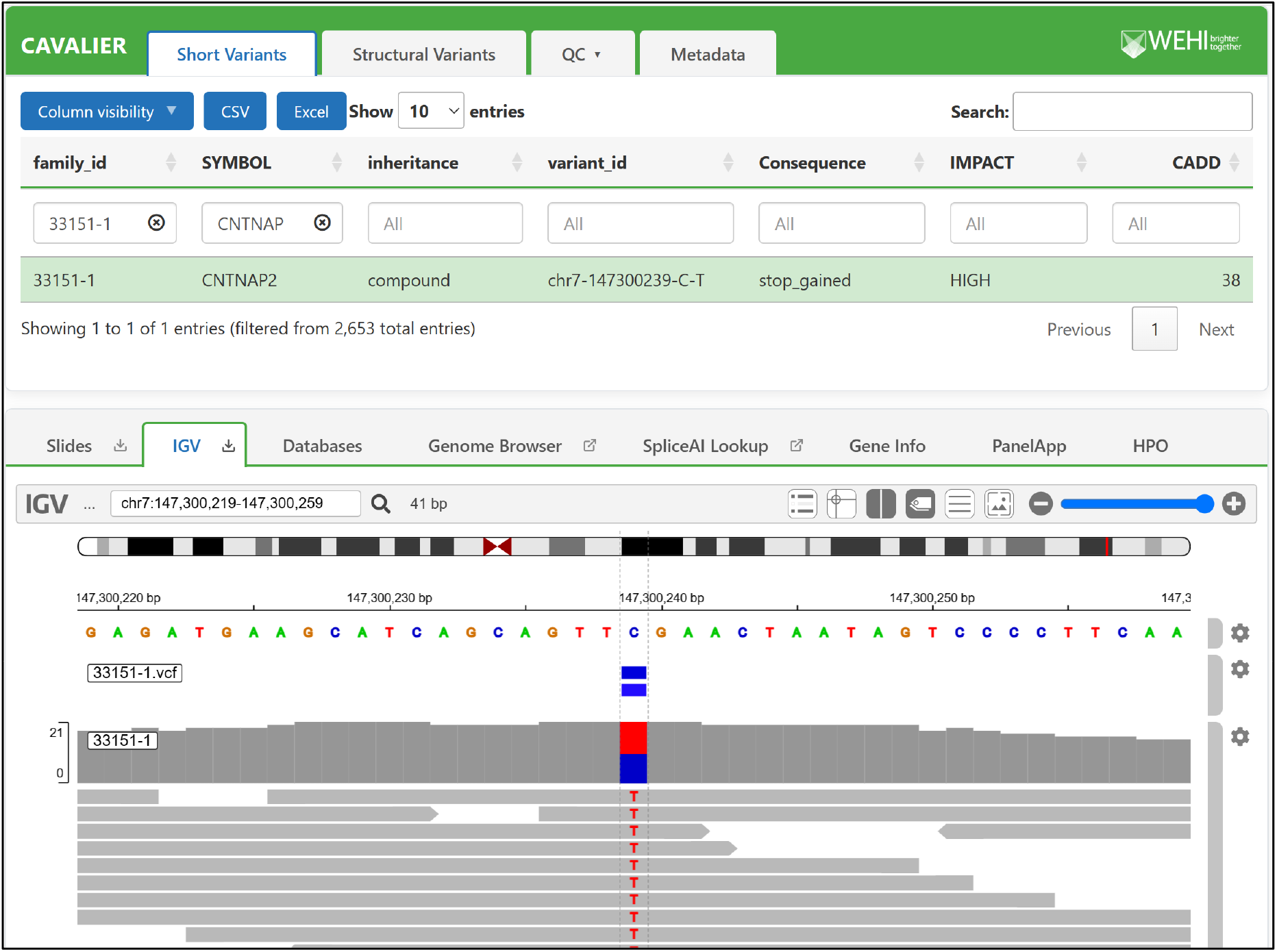
The CAVALIER interactive variant browser interface. The upper panel allows filtering of short and structural variants using customisable columns and includes tabs for QC data and run metadata. The lower panel enables exploration of variants and genes, with igv-reports (pictured) and other external APIs embedded.

In addition to the annotation, filtering, and reporting stages, the pipeline includes a QC workflow that runs somalier^22^ and SCE-VCF^23^ to assess relatedness, sex concordance, genetic ancestry, and contamination, enabling users to identify problematic samples and verify pedigree relationships.

## Research impact statement

**nf-cavalier** has been used in several studies investigating known and novel causes of rare disease in large-scale genome and exome sequencing cohorts, benefiting from extensive real-world application testing over five years. These include the Austin Health Adult Undiagnosed Disease Program (AHA-UDP), the first adult-focused undiagnosed disease program in Australia, which resulted in diagnoses for 16/50 (32%) participating families^24^. More recently, **nf-cavalier** has been used to search for molecular causes in a large cohort of individuals with developmental and epileptic encephalopathy (DEE), resulting in diagnoses for 37/242 (15%) of participants, including nine structural variants^21^, and in the identification of a complex structural variant in FBRSL1 associated with a severe DEE^25^. Furthermore, **nf-cavalier** was used to assist in the landmark discovery of an intronic repeat expansion in FGF14 as the most common cause of late-onset cerebellar ataxia, where it was used to exclude alternative explanations for pathogenicity^26^. The CAVALIER R package, a precursor to the current Nextflow implementation, was also used similarly to assist with the discovery of the RFC1 repeat expansion causing cerebellar ataxia with neuropathy and bilateral vestibular areflexia syndrome (CANVAS)^27^.

## AI usage disclosure

The majority of the **nf-cavalier** source code was written and tested without generative AI assistance. Since March 5, 2026 (GitHub commit ‘a1196d9’), Claude Code has been used for coding and documentation assistance, using a mixture of Sonnet 4.6 and Opus 4.7-4.8 models depending on task complexity. AI assistance was most useful for creating the interactive HTML output, updating and expanding the documentation, and implementing the pipelines to download the reference data and the test dataset. The AI tools were given clear, specific instructions on implementation and design, and all AI-generated content has been reviewed and tested by the authors. The authors made key decisions and take full responsibility for the software’s reliability and maintenance. The Claude web interface has also been used to assist with grammar, clarity, and phrasing for this manuscript.

## Acknowledgements

We thank Neblina Sikta (ORCID: 0000-0001-7778-805X) for contributions to the **nf-cavalier** GitHub repository. We would also like to thank members of the Epilepsy Research Centre (University of Melbourne, Austin Health) for their feedback and suggestions on the report output formats.

This research was supported by the Commonwealth through an Australian Government Research Training Program Scholarship (DOI: https://doi.org/10.82133/C42F-K220). This work was made possible through the Victorian State Government Operational Infrastructure Support Program and the Australian Government NHMRC IRIISS.

MB was funded by an Australian National Health and Medical Research Council Investigator grant (APP1195236). MFB was supported by an MRFF Genomics Health Futures Mission Grant (2007707). JEM was supported by a WEHI-CSL translational data science PhD top-up scholarship.

